# Environmental correlates of internal coloration in frogs vary throughout space and lineages

**DOI:** 10.1101/105684

**Authors:** Lilian Franco-Belussi, Diogo Borges Provete, Classius de Oliveira

## Abstract

Internal organs of ectotherms have melanin-containing cells. Several studies analyzed their developmental origin, role in immunity, and hormonal regulation. However, little is known about how environmental variables influence the distribution and quantity of organ coloration. Here, we addressed how environmental variables (temperature, UV, and photoperiod) influence the internal coloration of amphibians after controlling for spatial and phylogenetic autocorrelations. Coloration in all organs was correlated with phylogeny. However, the coloration of the heart, kidneys, and rectum of hylids, *R. schneideri,* some *Leptodactylus,* and *Proceratophrys* were influenced by temperature and photoperiod, whereas that of the testicle, lumbar parietal peritoneum, lungs, and mesenterium of Leiuperinae, Hylodidae, *Adenomera,* most *Leptodactylus* were influenced by UVB and temperature variation. Therefore, the amount of internal melanin seems to be a key trait influencing species distribution of frogs throughout space, since it can protect internal organs against the deleterious effect of high UV-B, temperature variation, and photoperiod.

**Significance:** The functions of internal coloration in fishes and frogs are little known. Internal pigmentation is commonly altered in fish and the degree of response is correlated with body transparency levels, suggesting possible adaptive functions. Here, we assume that internal melanin has protective functions against UV-B, temperature variation, and photoperiod. Thus it could influence frogs species distribution throughout space. The melanin coloration of each organ was influenced by distinct environmental variables depending on the lineages of species. Our results could direct further studies about the functions of internal coloration.

## Introduction

Vertebrates have a variety of body color patterns whose evolution is shaped by both natural and sexual selection (Aspengren et al., 2009). The body color of ectotherms may vary in response to environmental changes in a number of ways. The mechanisms underlying those changes include physiological color changes that works by a rapid pigment transportation in the cell cytoplasm (Bagnara and Matsumoto, 2006; Aspengren et al., 2009; Sköld et al., 2012). Cells responsible for color changes in fish and amphibians are called chromatophores, which can be subdivided into melanophores, erythrophores, xanthophors, and iridophores (Wallin, 2002; Bagnara and Matsumoto, 2006).

The internal organs and structures of ectotherms can also have different color patterns, which are given by different pigments. One of these pigments is melanin, which occurs in cells called melanocytes. These cells produce and store melanin (Agius and Roberts, 2003; Oliveira and Franco-Belussi, 2012) and are similar to skin melanocytes (Zuasti et al., 1998; Franco-Belussi et al., 2013). Melanocytes occur in internal organs and membranes of amphibians (Oliveira and Franco-Belussi, 2012) and fish (Agius, 1981).

Environmental variables influence physiological and behavioral aspects of ectotherms. For example, temperature can alter the internal coloration in anurans, due to the thermoprotective role of melanin-containing cells. High temperature decreases liver pigmentation in anurans (Santos et al., 2014). At low temperatures, animal’s metabolism may be reduced. As a consequence, melanin in the liver increases (Barni et al., 1999).

Ultra-Violet (UV) radiation is one of the factors responsible for amphibian declines (Blaustein et al., 1997). Exposure to UV radiation affects embryonic development, causing morphological changes and tadpole mortality, which in turn result in population declines (Lipinski et al., 2016). Internal melanin seems to protect internal organs against the genotoxic effects of UV radiation (Roulin, 2014). For example, short-term (e.g., 24 h) exposure to UV radiation increases the coloration in both skin and organ’s surface in adult anurans (Franco-Belussi et al., 2016).

Another environmental variable that alters the skin coloration of fish and anurans is photoperiod. Light disperses melanin granules in cells of amphibians, increasing coloration (Moriya et al., 1996). Previous studies showed that light intensity and time of exposure increase the skin color of fish (Han et al., 2005). Also, the secretion of Melanin Concentrating Hormone (MCH) increases in fish after long photoperiods (Lyon and Baker 1993). The MCH is involved in the aggregation of melanin granules, which makes the animal lighter. However, little is known about how environmental factors influence the coloration of visceral organs in anurans.

The effects of environmental factors on internal coloration were analyzed under experimental conditions and in a few species. The main goal of these experimental studies was to demonstrate the mechanistic basis of the responses of pigmentary cells to these factors. The only large-scale study that evaluated the distribution of coloration categories on organs of 32 Neotropical anuran species (Provete et al., 2012) found that the intensity of coloration in several organs has a phylogenetic component. Nonetheless, little is known about how the organ’s coloration of anuran species vary in response to environmental factors over broad spatial scales.

Here, we expand our previous study (Provete et al., 2012) to address this question using multivariate methods (Pavoine et al., 2011) considering intraspecific variation in organ coloration. Specifically, we asked how environmental variables (temperature, UV, and photoperiod) influence the internal coloration of anuran species after controlling for spatial and phylogenetic autocorrelations. Internal melanin can possibly confer adaptive advantages for species in response to environmental variables, as mentioned above. Therefore, internal melanin may influence species distribution throughout space.

## Results

### Phylogenetic pattern of organ coloration

Coloration considering all organs was correlated with the phylogeny. The greatest diversity was found in Leptodactylidae (Figure 1), followed by the node that includes all species except for Brachycephalidae and Microhylidae. The highest level of intraspecific variation occurred in *D. minutus* and to a lesser extent in Dendropsophryni (Figure 1). The organs that had the greatest diversity of coloration were the testis, heart, rectum, mesentery, peritoneum, and lung (Figures S1-S7).

**Figure 1.**
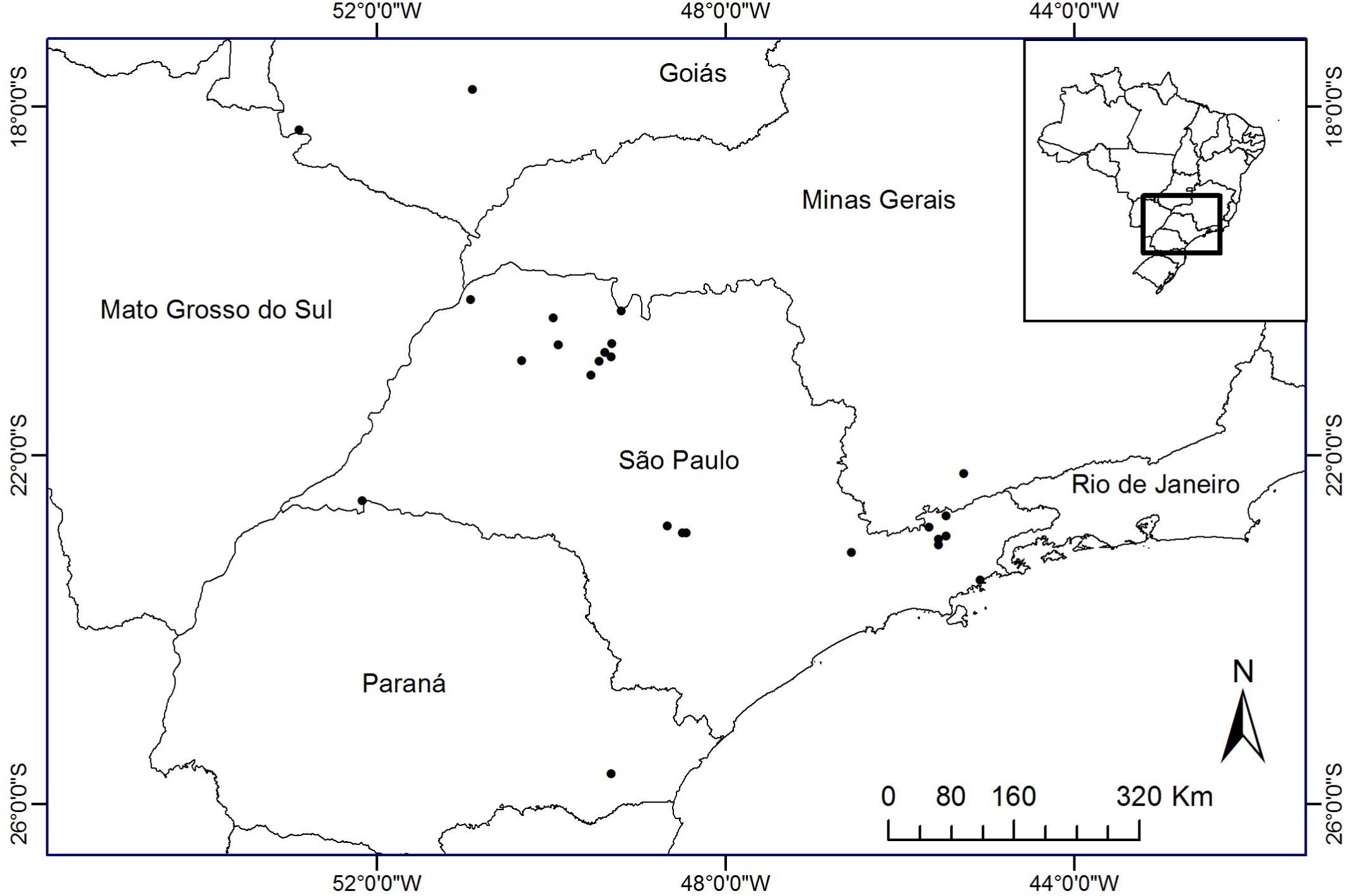
Decomposition of diversity of organ coloration along the nodes of the phylogeny considering all organs together. For species names see Figure 7. The high values of diversity are skewed towards the root of the phylogeny, indicating phylogenetic signal.

### Relationships between organ coloration and environmental variables

The co-inertia of first axis of the RLQ is 0.67, which is equivalent to 67% of the total variation. The positive side of the first axis corresponds to areas with high photoperiod. The localities most positively correlated with the first ordination axis were Ubatuba, Fazenda Rio Grande, Atibaia, Taubaté, and Rio Verde (Figure 2). Therefore, these localities had a high Bio10 and photoperiod (Figure 3A). The species found in those localities have similar coloration categories in the heart, kidneys, and rectum (Figure 3B). These species are most hylids, *R. schneideri,* some *Leptodactylus,* and *Proceratophrys* (Figure 4).

**Figure 2.**
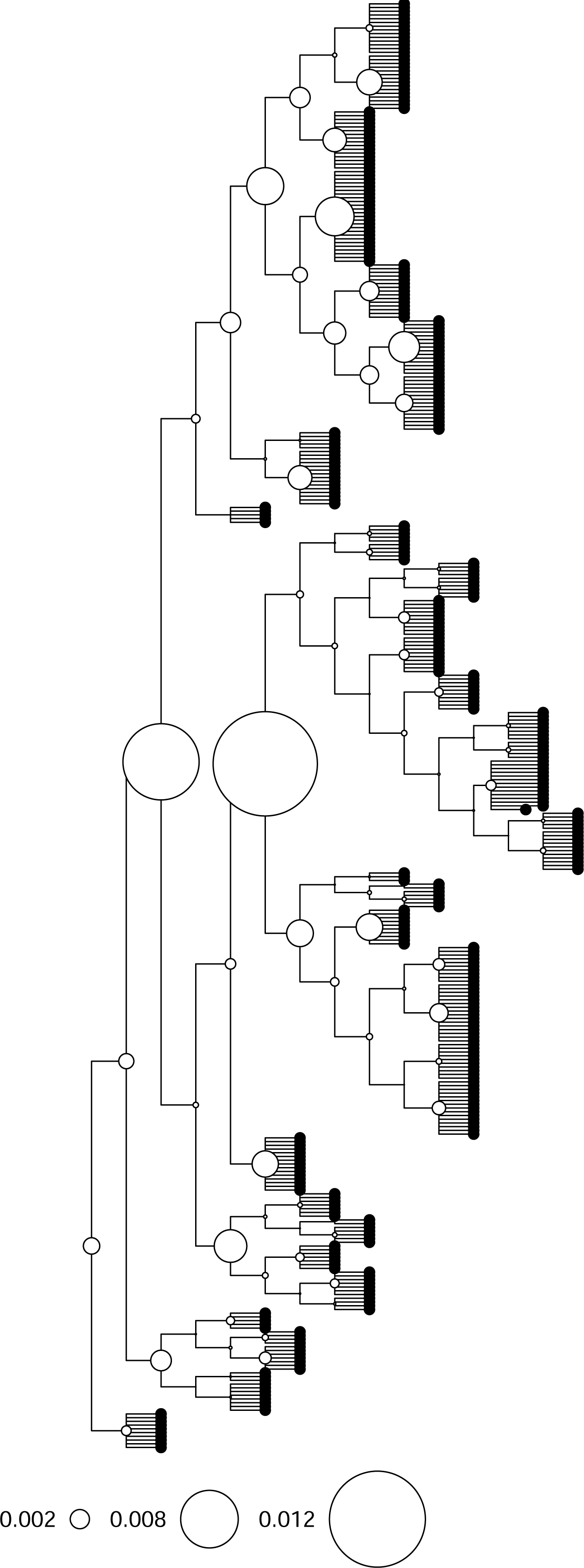
Results of the RLQ analysis visualized in space. Circles show the localities. The coordinates of the sites were analyzed respective to the first ordination axis. Sites in black are positively correlated to the first axis, while blank sites were negatively correlated. Size of circles indicate the absolute value of the coordinates.

**Figure 3.**
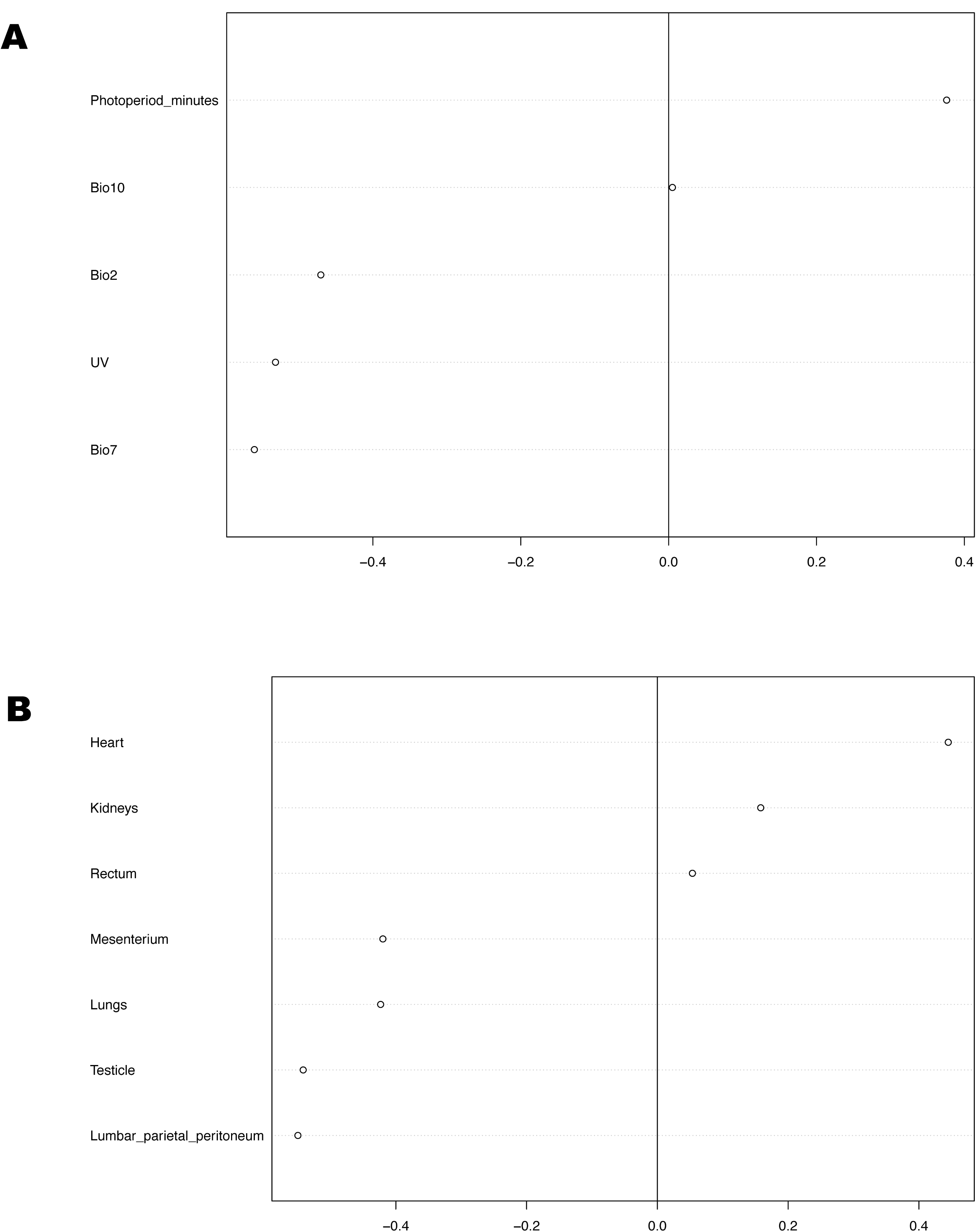
Effects of environmental variables and organs on the first axis of RLQ. A) Spearman rank-correlations between organ coloration category (ordinal) and the coordinates of species on the first axis. B) Pearson correlation between the environmental variables and the coordinates of sites in the first axis. Species with high coloration on the heart, kidney, and rectum occur in sites with high BIO10 and photoperiod, whereas those with high coloration on the mesenterium, lungs, testicle, and peritoneum occurred in sites with high BIO2, BIO7, and UVB.

**Figure 4.**
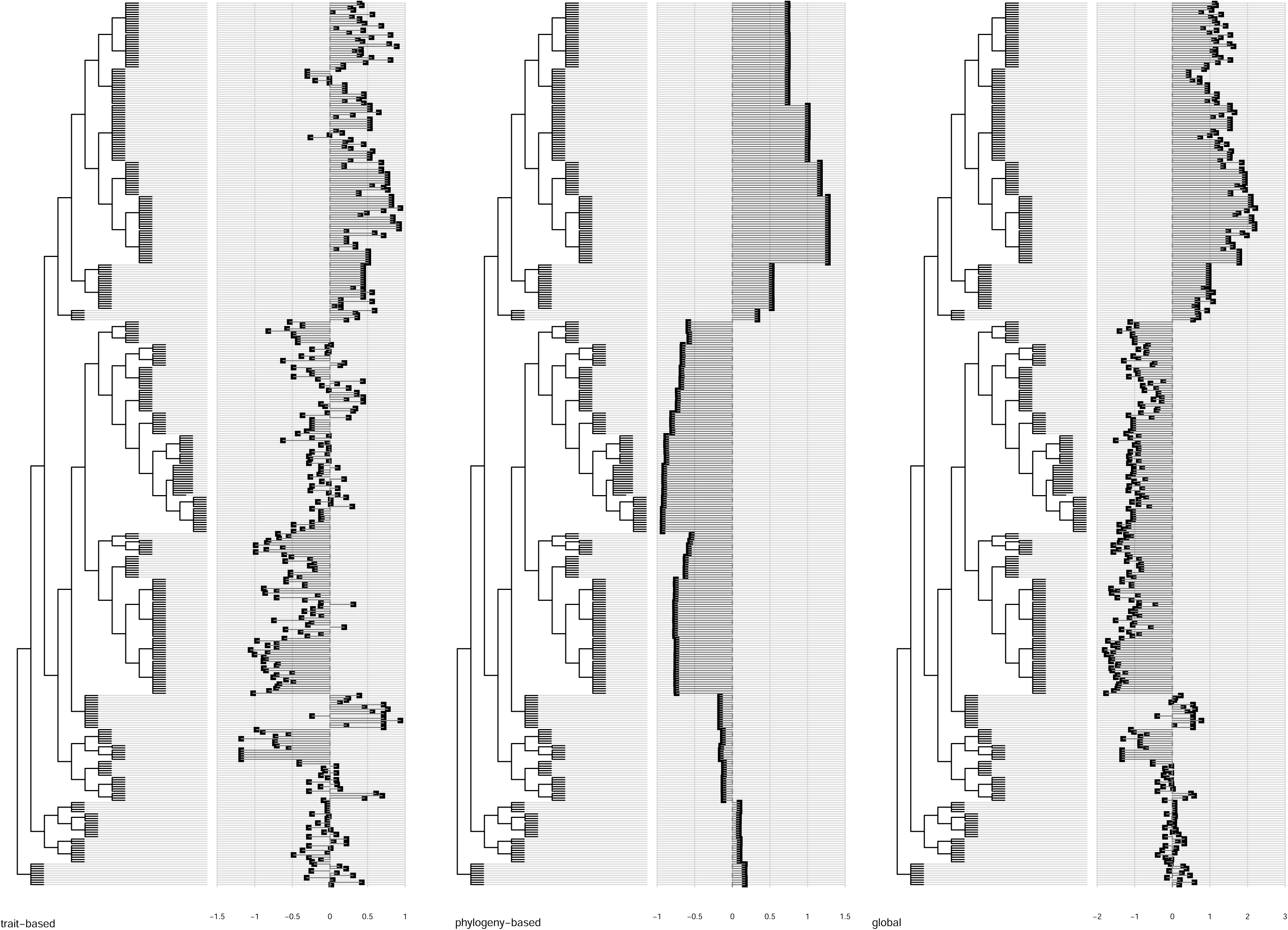
Results of the RLQ analysis (first axis) visualized on the phylogeny. The coordinates of the species are the sum of combination of trait and phylogenetic variables.

The negative side of the axis 1 represents localities with high UVB, Bio2, and Bio7 (Figure 3A). The localities most negatively correlated with the first axis are Cristina and those in the Northwest of the state of São Paulo, such as Santa Fé do Sul, Icém, Santo Antonio do Aracanguá, Teodoro Sampaio, Votuporanga, and Nova Itapirema (Figure 2). The species found in these localities have similar coloration categories in the testicle, lumbar parietal peritoneum, lungs, and mesenterium (Figure 3B). The species are Leiuperinae, Hylodidae, *Adenomera,* most *Leptodactylus,* and to a lesser extent the Brachycephaloidea and *Elachistocleis* (Figure 4).

## Discussion

We found that the internal coloration in anurans was influenced by environmental variables and has a non-stationary phylogenetic component, mainly in the testicle and peritoneum. Interestingly, we also found a large intraspecific variation in coloration intensity in some lineages, mainly the Dendropsophryni tribe. In addition, different organ coloration appears to be influenced by different environmental variables. Interestingly, the mean value of organ coloration did not show a clear spatial pattern.

Both the diurnal (BIO2) and the annual (BIO7) thermal ranges influenced more strongly the coloration in the testicle, lumbar parietal peritoneum, lungs, and mesenterium. Melanin-containing cells protect tissues against temperature variation. Melanin decreases in hibernating species during the winter, due to both decreased synthesis and cellular apoptosis (Barni et al., 2002). Similarly, temperature variation reduced the hepatic pigmentation in a Neotropical anuran species (Santos et al., 2014), demonstrating that temperature is a key environmental factor controlling the amount of melanin in the liver. Melanin can also dissipate heat and is probably involved in thermoregulation in ectotherms (Cesarini, 1996). Thus, the amount of melanin may be a trait with possible adaptive functions that determine the occurrence of species in localities with wide variation in temperature.

UV-B radiation influences coloration in the same organs as temperature range. A previous study found that short-term exposure to low doses of UV-B increases internal coloration by increasing melanin production and dispersion (Franco-Belussi et al., 2016). UV radiation can cause genotoxic effects in cells by disrupting DNA (Ortonne et al., 2002). Melanin provides protection against solar radiation, by dissipating solar energy in the form of heat (Ortonne et al., 2002). As a result, tyrosinase activity increases in melanin-containing cells, which increases melanin production to protect cells against UV radiation (Friedmann and Gilchrest, 1987). Therefore, species that occur in sites with high UV-B incidence tend to have greater production of melanin as a mechanism to deal with its deleterious effects.

Photoperiod influenced the coloration on the heart, kidneys, and rectum. The effects of photoperiod on internal coloration are poorly known. However, photoperiod directly influences the endocrine system (Breet, 1979) that can indirectly alter melanin amount. An increase in photoperiod (e.g., 18:6 light:dark) promotes whitening of fish’s skin by increasing the secretion of MCH (Lyon and Baker, 1993; Guinés et al., 2004). The adaptive value of coloration on the heart, kidneys, and rectum is still not clearly understood (Colombo et al., 2011), but it is probably related to the functions of the melanin molecule which mainly acts as antibiotic, in light absorption (e.g., photoprotection), cation chelator, and free radical sink (Riley, 1997). Additionally, the intensity of coloration on the heart and kidneys of hylids tends to be lower than in the testicles of Leiuperinae, showing a phylogenetic signal.

We found that the amount of melanin on a given organ is determined jointly by its physiology, environmental variables, and phylogenetic relationship (see also Provete et al., 2012). Melanocytes have distinct physiology depending on the external coloration of the animal. For example, pigmented cells on the peritoneum of fish respond to hormones, such as melatonin and epinephrine (Sköld et al., 2010). However, the aggregation or dispersion of pigmented cells promoted by the hormone depends on the transparency of the animal (Sköld et al., 2010). This demonstrates that the internal pigment cells can adapt to distinct situations, behaving differently in animals depending on their cutaneous coloration (Sköld et al., 2010). In addition, changes in internal color in transparent animals may be related to substrate adaptation or social signaling (Sköld et al., 2010). These results reinforce the role of physiological responses of pigmented cells.

As UV radiation and temperature can have deleterious effects, species that occur in places with high incidence of these factors could have developed more melanin on the testicles as to protect their germinal epithelium, since damages in the gametes can influence the reproductive fitness of individuals. For example, hylodids have a large amount of melanin on the testicles and are restricted to the Atlantic rainforest. This region has the same degree of UV incidence of the northwest of São Paulo, where swamp frogs of the subfamily Leiuperinae occur. As a consequence, these species developed similar strategies to deal with elevated UV-B variation by having high amount of coloration on the testicles. Also, having a high amount of melanin on the testicles may allow species to be active during the day, such as dendrobatids (Grant et al., 2006), or at dusk like *Pseudopaludicola* and some *Physalaemus* (Vasconcelos and Rossa-Feres, 2005). Conversely, species lacking melanin on the testicles are mainly active at night (e.g., Hylidae and Leptodactylidae; Vasconcelos and Rossa-Feres, 2005).

Therefore, the amount of internal melanin seems to be a key trait influencing anuran species distribution throughout space, since it can protect internal organs against the deleterious effect of high UV-B, temperature variation, and photoperiod.

## Methods

### Specimen sampling

The anuran species used in this study were collected at night when calling, near breeding sites in 26 localities in the states of São Paulo and Goiás, which are housed at the collection of the Laboratório de Anatomia - UNESP. We used at least five adult males of each species for the analysis of pigmentation. The specimens were anesthetized with 5 g/L of benzocaine and dissected to expose the organs. All procedures followed the recommendations of the COBEA (Brazilian College of Animal Experimentation) and the Ethics Committee of our university (Protocol #70/07 CEEA). We also analyzed additional specimens from the amphibian collections of the Department of Zoology and Botany, UNESP (DZSJRP); Scientific Collection of the Laboratory of Zoology, University of Taubaté (CCLZU); and the Jorge Jim collection (JJ; now incorporated to the herpetological collection of the National Museun, MN-RJ) from three localities in the states of Paraná and Minas Gerais (Figure 5). All material examined is listed in Appendix 1. We also obtained data from the literature (Franco-Belussi et al., 2011; Franco-Belussi et al., 2012) for species of the family Hylidae. In total, we had 388 specimens from 43 species belonging to six families. Species had different sample sizes because same of them were widely distributed. Thus, we wanted to asses if intraspecific variation in coloration was somehow related to spatial variation in coloration.

**Figure 5.**
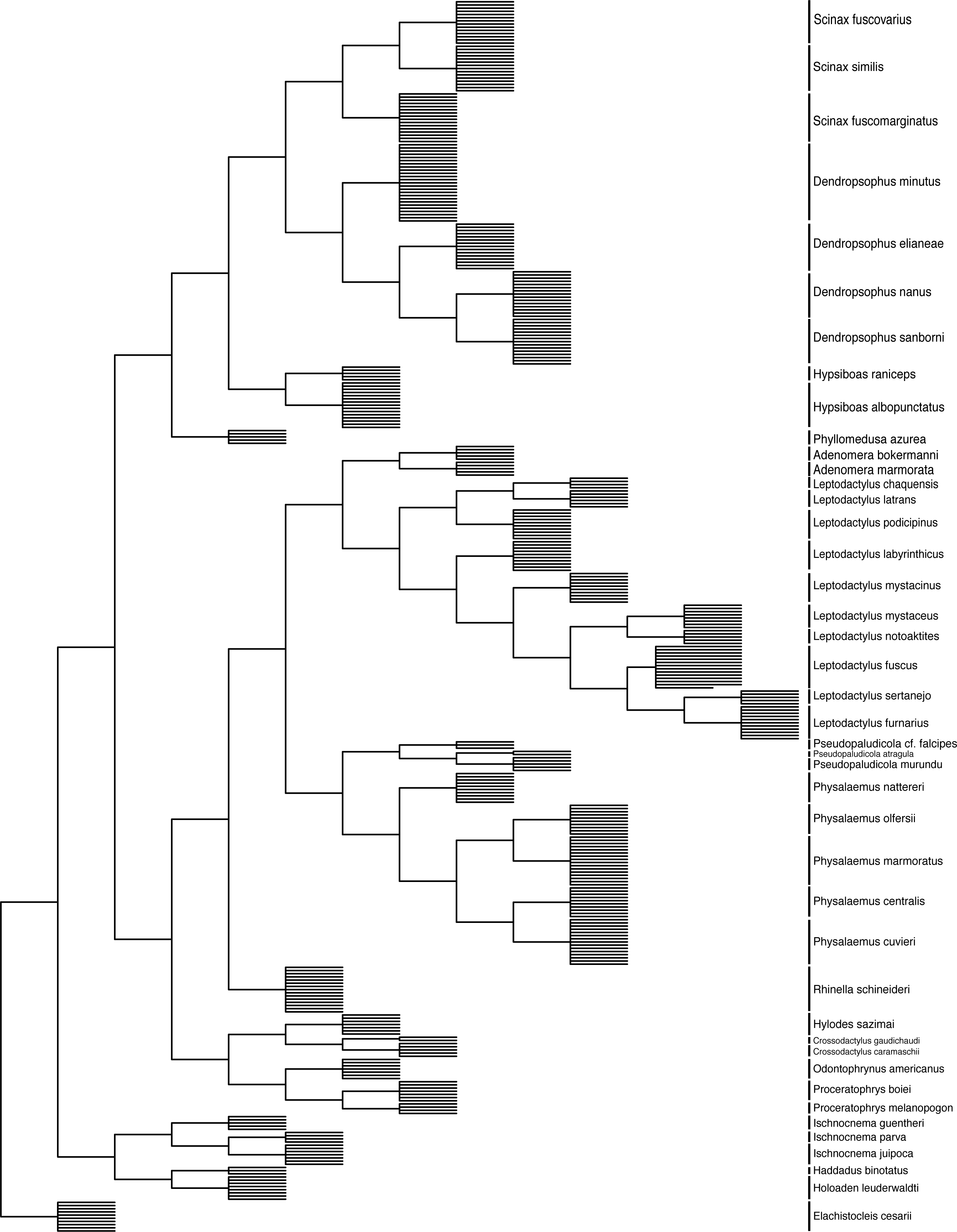
Map showing the sites sampled in this study and those to which we obtained data from the literature or museum.

### Morphological data

We recorded the distribution of visceral melanocytes in 15 organs or structures using a Leica stereoscopic microscope (MZ16), coupled with an image capture system, namely: heart, lungs, rectum, peritoneum, kidneys, testes, and intestinal mesenterium. For each individual, we recorded the coloration on these organs/structures based on coloration intensity, following the protocol of Franco-Belussi et al. (2009). Briefly, the intensity of organ coloration was divided into four categories, ranging from absence to entirely colored, as follows: Category 0) absence of pigment cells on the surface of organs, in which the usual color of the organ is evident; Category 1) a few scattered pigment cells, giving the organs a faint pigmentation; Category 2) a large amount of pigment cells; Category 3) a massive amount of pigment cells, rendering an intense pigmentation to the structure, changing its usual color and superficial vascularization (Franco-Belussi et al., 2009). Thus, for a given organ or structure, each individual could display four categories of pigmentation: 0, 1, 2, or 3. Pigmentation category was assessed in a double blind fashion. Pictures of organs and regions are freely available in Morphobank at http://dx.doi.org/10.7934/P701.

To minimize multicollinearity, we calculated the Variation Inflation Factor (VIF; Zuur et al., 2010) and a pair-wise correlation for the organs. Cardiac blood vessels, Renal veins, and Lumbar nerve plexus had a high VIF. Therefore, we excluded them from further analysis.

### Statistical analyses

We extracted the bioclimatic variables Bio2, Bio3, Bio4, Bio5, Bio6, Bio7, and Bio10 related to temperature from WorldClim (Hijmans et al., 2005) for the 26 localities. Data for photoperiod (minutes of light-hours in the rainy season, when most species were collected) were obtained from the Brazilian National Observatory (Brasil 2016)). Data for UV-B radiation were extracted from a raster file (Beckmann et al., 2014). We standardized all variables to zero mean and unit variance previously to analysis. Posteriorly, we tested for multicollinearity (Zuur et al., 2010) and removed environmental variables with VIF > 10. The reduced variables were Bio2, Bio7, Bio10, UVB, and photoperiod. We then tested for spatial correlation in the environmental variables. All environmental variables were spatially autocorrelated, with Moran’s *I* varying between 0.154 and 0.485. The reduced matrix of environmental variables was analyzed with a Principal Component Analysis (PCA; Legendre and Legendre, 2012).

The phylogeny for the species to which we had trait data was pruned from the dated phylogeny of Pyron (2014) for amphibians. This phylogeny was inferred based on nine nuclear genes and three mitochondrial genes for 3,309 species, with average 20% of completeness. To this topology, we added each individual as a polytomy to its corresponding species with branch length equal to unit (Figure 6). Since our trait is categorical, we could not simply calculate its standard error to account for intraspecific variation. Then, we extracted a distance matrix from this phylogeny and calculated a Principal Coordinates Analysis (PCoA; Legendre and Legendre, 2012) to extract phylogenetic eigenvectors.

**Figure 6.**
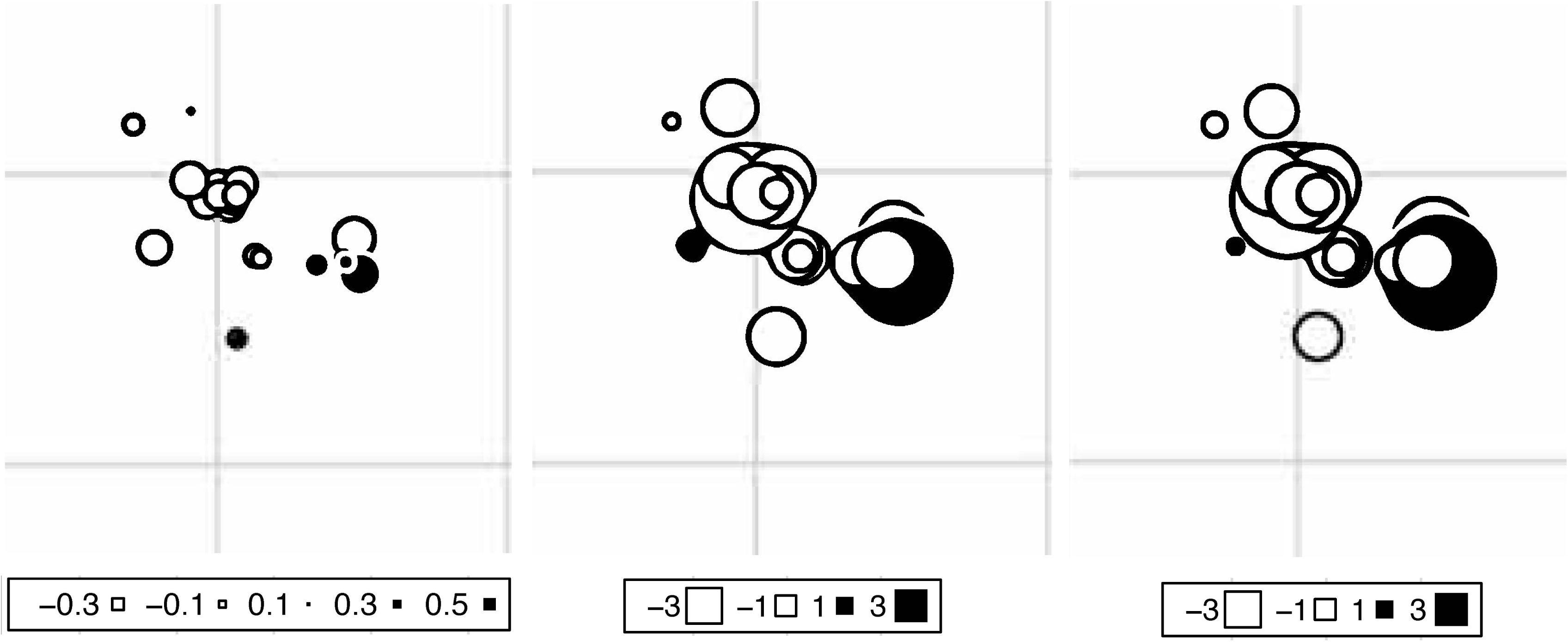
Phylogeny of 43 species from six families used in this study with all 388 individuals included as polytomy.

For the species composition matrix, the presence of each individual analyzed was placed in rows and localities as columns. This matrix was analyzed with a Correspondence Analysis (CA).

To model space, we built a neighbor matrix linking sites separated up to 318.88 Km (based on the truncation distance of a Minimum Spanning Tree). Then, we computed a PCA onto this neighbor matrix to use as spatial variables in the extended RLQ.

The trait matrix contained the coloration category for each individual (rows) in each organ (columns). We then tested for phylogenetic correlation (phylogenetic “signal”) in the coloration of each organ by decomposing the trait diversity, calculated as Rao’s entropy, along the nodes of the phylogeny (Pavoine et al., 2010). The Rao’s quadratic entropy only uses tree topology to decompose trait diversity. Afterwards, we tested if the diversity of coloration categories was biased towards the root of the phylogeny, or concentrated in a single or a few nodes (Pavoine et al., 2010). In this context, a phylogenetic signal occurs when trait diversity is skewed towards the root of the phylogeny, implying that all its descending lineages would have similar values for that trait. We found a phylogenetic signal in the coloration of internal organs when we consider them altogether (Table S1), but not separately. Then, we calculated a distance matrix for traits based on the modified Gower similarity coefficient (Pavoine et al., 2009). Posteriorly, we tested for a relationship between environmental variables and the coloration of each organ using a multivariate version of the Fourth-corner analysis. Significance was tested using the null model 4 (Dray and Legendre, 2008). All organs, but the pericardium had significant relationship with environmental variables. Thus, we excluded this organ from further analysis. Then, we calculated an Euclidian distance matrix for the categories of organ coloration (ordinal data). This distance matrix was then analyzed with PCoA. Finally, we used an extended version of the RLQ ordination (Pavoine et al., 2011) that takes into account the spatial dependency of environmental variables and the phylogenetic autocorrelation in species traits to test the influence of environmental variables on species traits. Analyses were implemented in R v. 3.3.2 (R Core Team 2016) package ade4 (Dray and Dufour, 2007) and functions provided by Pavoine et al. (2011). Data and an R script used to conduct analyses are available in FigShare (Franco-Belussi et al., 2017).

## Acknowledgments

The study was supported by a research grant from the São Paulo Research Foundation - FAPESP (#2015/12006-9 and 05/02919-5) to CO. and postdoc fellowships to LFB (#2014/00946-4) and DBP (#2016/13949-7). LFB was also supported by a CAPES-PNPD postdoc fellowship during the final preparation of this manuscript. T. Gonfalves-Souza and F. R. da Silva helped interpreting the analysis output. L. R. S. Santos, I. A. Martins, S. A. César, R. M. Moresco, and R. Zieri helped with specimen sampling. L. R. S. Santos provided additional specimens. B. Vilela and D. P. Silva kindly prepared the map. *Authors’ contributions:* LFB collected the data. DBP analyzed and curated the data, prepared the figures and supplementary material. LFB and DBP prepared the first draft of the manuscript. CO designed the study, contributed chemicals and equipment. All authors read and approved the final manuscript.

## Supplementary material

Figure S1. Decomposition of the diversity of organ coloration along the nodes of the phylogeny. A) Testicle; B) Rectum; C) Heart; D) Lungs; E) Kidneys; F) Peritoneum; G) Mesenterium.

Table S1. Results of the phylogenetic correlation analysis for each organ separately and all together.

## Appendix 1.

List of all specimens analyzed.

## References

Agius, C. (1981). Preliminary studies on the ontogeny of the melanomacrophages of teleost hematopoetic tissues and age-related changes. Dev. Comp. Immunol. 5, 597–606

Agius, C. and Roberts, R.J. (2003). Review: Melano-Macrophage Centres and their Role in Fish Patology. J. Fish Biol. 26, 499–509

Aspengren, S., Hedberg, D., Sköld, H.N. and Wallin M. (2009). New Insights into Melanossome Transport in Vertebrate Pigment Cells. Int. Rev. Cell Mol. Biol. 272, 245–302

Bagnara, J.T. and Matsumoto, J. (2006). Comparative Anatomy and Physiology of Pigment Cells in Nonmammalian Tissues. In: The Pigmentary System: Physiology and Pathophysiology, Nordlund J. J., Boissy, R.E., Hearing, V.J., King, R. A., Ortonne, J.P. eds. (New York, Oxford: Oxford University Press) pp.9–40

Barni, S., Vaccarone R., Bertone V., Fraschini A., Bernini F. and Fenoglio C. (2002). Mechanisms of changes to the liver pigmentary componente during the annual cycle (activity and hibernation) of *Rana esculenta* L. J. Anat. 200,185–194

Barni, S., Bertone, V., Croce, A.C., Bottirolli, G., Bernini, F. and Gerzeli, G. (1999). Increase in Liver Pigmentation During Natural Hibernation in some Amphibians. J. Anat. 195,19–25

Beckmann, M.T., Vaclavik, A.M., Manceur, L., Šprtová, H., von Wehrden, E., Welk, A. F., Cord, and Tatem, A. (2014). glUV: a global UV-B radiation data set for macroecological studies. Methods Ecol. Evol. 5, 372–383

Blaustein, A. R., Kiesecker, J. M., Chivers, D. P., and Anthony, R. G. (1997). Ambient UV-B radiation causes deformities in amphibian embryos. Proc. Nat. Acad. Sc. 94, 13735–13737

Brasil. (2016). Anuário do observatório nacional - efemérides astronômicas. Rio de Janeiro: Observatório Nacional - Ministério da Ciência, Tecnologia e inovação.

Brett, J.R. (1979). Environmental factors and growth. In: Fish Physiology, Vol. VIII, W.S. Hoar, D.J. Randall and J.R. Brett eds., (Academic Press, New York, USA) pp. 599–675

Cesarini, J.P. (1996). Melanins and their possible roles through biological evolution. Adv Space Res. 18, 35–40

Colombo, S., Berlim, I., Delmas, V. and Larue, L. (2011). Classical and non-classical melanocytes in vertebrates. In: Melanins and Melanosomes: Biosynthesis, Biogenesis, Physiological and Pathological Functions, 1st ed. Borovanský J, Riley PA eds., pp. 21–62

Dray, S. and Dufour, A.B. (2007). The ade4 package: implementing the duality diagram for ecologists. J. Stat. Softw. 22, 1–20

Dray, S. and Legendre, P. (2008). Testing the species traits-environment relationships: the fourth-corner problem revisited. Ecology. 89,3400–3412

Franco-Belussi, L., Zieri, R., Santos, L.R.S., Moresco, R.M. and Oliveira, C. (2009). Pigmentation in Anuran Testes: Anatomical Pattern and Variation. Anat. Rec. 292,178182

Franco-Belussi, L., Santos, L. R. S., Zieri, R., and Oliveira, C. (2011). Visceral pigmentation in four *Dendropsophus species* (Anura: Hylidae): Occurrence and comparison. Zool. Anz. 250,102–110

Franco-Belussi, L., Santos, L. R. S., Zieri, R., and Oliveira, C. (2012). Visceral pigmentation in three species of the genus *Scinax* (Anura: Hylidae): distinct morphological pattern. Anat. Rec. 295, 298–306

Franco-Belussi, L., Castrucci, A.M.D.L. and Oliveira, C. (2013). Responses of melanocytes and melanomacrophages of *Eupemphix nattereri* (Anura: Leiuperidae) to Nle^4^, D-Phe-a-melanocyte stimulating hormone and lipopolysaccharides. Zool. 116, 316–324.

Franco-Belussi, L., Sköld, H. N. and Oliveira, C. (2016). Internal pigment cells respond to external UV radiation in frogs. J. Exp. Biol. 219, 1378–1383

Franco-Belussi L, Provete D. and Oliveira C. (2017). Environmental correlates of internal coloration in anurans vary throughout space and lineages. Figshare https://doi.org/10.6084/m9.figshare.4707187.v1

Friedmann, P. S. and Gilchrest, B. A. (1987). Ultraviolet radiation directly induces pigment production by cultured human melanocytes. J. Cell. Physiol. 133, 88–94

Ginés, R., Afonso, J.M., Argüello, A., Zamorano, M.J., and López, J. L. (2004). The effects of long-day photoperiod on growth, body composition and skin colour in immature gilthead sea bream *(Sparus aurata* L.). Aquac. Res. 35, 1207–1212

Grant, T., Frost, D.R., Caldwell, J. P., Gagliardo, R., Haddad, C.F.B., Kok, P. J. R., Means, D. B., Noonan, B. P., Schargel, W. E. and Wheeler, W. C. (2006). Phylogenetic systematics of the dart-poison frogs and their relatives (Amphibia: Athesphatanura: Dendrobatidae). Bull. Am. Mus. Nat. Hist. 1–262.

Han, D., Xie, S., Lei, W., Zhu, X., and Yang, Y. (2005). Effect of light intensity on growth, survival and skin color of juvenile Chinese longsnout catfish (*Leiocassis longirostris* Günther). Aquac. 248, 299–306

Hijmans, R. J., Cameron, S.E., Parra, J. L., Jones, P. G. and Jarvis, A. (2005). Very high resolution interpolated climate surfaces for global land areas. Int. J. Climatol. 25, 1965–1978

Legendre, P., Legendre, L. (2012). Numerical Ecology. 3rd. edition. Elsevier Limited, Oxford.

Lipinski, V.M., Santos, T. G. and Schuch, A. P. (2016). An UV-sensitive anuran species as an indicator of environmental quality of the Southern Atlantic Rainforest. J. Photochem. Photobiol. B: Biol. 165,174–181.

Lyon, V.E. and Baker, B.I. (1993). The effect of photoperiod on plasma levels of melaninconcentrating hormone in the trout. J. Neuroendocrinol. 5, 493–499

Moriya, T., Miyashita, Y., Arai, J. I., Kusunoki, S., Abe, M., and Asami, K. (1996). Light-sensitive response in melanophores of *Xenopus laevis:* I. Spectral characteristics of melanophore response in isolated tail fin of *Xenopus* tadpole. J. Exp. Zool. 276, 11–18

Oliveira, C. and Franco-Belussi, L. (2012). Melanic pigmentation in ectothermic vertebrates: occurrence and function. In: Melanin: Biosynthesis, Functions and Health Effects, Nova Publishers, pp. 213–226.

Ortonne, J.P. (2002). Photoprotective properties of skin melanin. Br. J. Dermatol. 146, 7–10.

Pavoine, S., Vallet, J., Dufour, A.-B., Gachet, S., and Daniel, H. (2009). On the challenge of treating various types of variables: application for improving the measurement of functional diversity. Oikos 118, 391–402.

Pavoine, S., Baguette, M., and Bonsall, M. B. (2010). Decomposition of trait diversity among the nodes of a phylogenetic tree. Ecol. Monog. 80, 485–507.

Pavoine, S., Vela, E., Gachet, S., Belair, G. and Bonsall, M.B. (2011). Linking patterns in phylogeny, traits, abiotic variables and space: a novel approach to linking environmental filtering and plant community assembly. J. Ecol. 99, 165–175

Provete, D.B., Franco-Belussi, L., Santos, L.R.S., Zieri, R., Moresco, R.M., Martins, I. A., Almeida, S.C. and Oliveira, C. (2012). Phylogenetic signal and variation of visceral pigmentation in eight anuran families. Zool. Scripta, 41, 547–556

Pyron, R.A. (2014). Biogeographic analysis reveals ancient continental vicariance and recent oceanic dispersal in amphibians. Syst. Biol. 63, 779–797

R Core Team. (2016). R: A language and environment for statistical computing. R Foundation for Statistical Computing, Vienna, Austria. URL https://www.R-project.org/

Riley, P.A. (1997). Melanin. Int. J. Biochem. Cell Biol. 29, 1235–1239

Roulin, A. (2014). Melanin-based colour polymorphism responding to climate change. Global Change Biol. 20, 3344–3350

Santos, L.R.S., Franco-Belussi, L., Zieri, R., Borges, R.E. and Oliveira, C. (2014). Effects of Thermal Stress on Hepatic Melanomacrophages of *Eupemphix nattereri* (Anura). Anat. Rec. 297, 864–875

Sköld, H.N., Aspengren S. and Wallin M. (2012). Rapid color change in fish and amphibians - function, regulation, and emerging applications. Pig. Cell Melan. Res. 26, 28–39

Sköld, H.N., Svensson, P.A. and Zejlon, C. (2010). The capacity for internal colour change is related tobody transparency in fishes. Pig. Cell Melan. Res. 23, 292–295

Vasconcelos, T. S. and D. C. Rossa-Feres. (2005). Diversidade, distribuição espacial e temporal de anfíbios anuros (Amphibia, Anura) na região noroeste do estado de são paulo, brasil. Biota Neotropica, 5,1–14

Zuasti, A., Jiménez-Cervantes, C., García-Borrón, J.C., Ferrer, C. (1998). The melanogenic system of *Xenopus laevis.* Arch. Histol. Cytol. 61, 305–316

Zuur, A. F., Ieno, E.N. and Elphick, C.S. (2010). A protocol for data exploration to avoid common statistical problems. Methods Ecol. Evol. 1, 3–14

Wallin, M. (2002). Nature’s Palette: How Animals, Including Humans, Produce Colours. Biosc. Explain. 1,1–12

